# Evolution of sex-biased genes in *Drosophila* species with neo-sex chromosomes: potential contribution to reducing sexual conflict

**DOI:** 10.1101/2023.10.28.564560

**Authors:** Anika Minovic, Masafumi Nozawa

## Abstract

An advantage of sex chromosomes is the potential to reduce sexual conflict because they provide a basis for selection to operate separately on females and males. However, evaluating the relationship between sex chromosomes and sexual conflict is challenging owing to the difficulty in measuring sexual conflict and substantial divergence between species with and without sex chromosomes. We therefore examined sex-biased gene expression as a proxy for sexual conflict in three sets of *Drosophila* species with and without young sex chromosomes, the so-called neo-sex chromosomes. In all sets, we detected more sex-biased genes in the species with neo-sex chromosomes than in the species without neo-sex chromosomes in larvae, pupae, and adult somatic tissues but not in gonads. In particular, many unbiased genes became either female- or male-biased after linkage to the neo-sex chromosomes in larvae, despite the low sexual dimorphism in larvae. However, sexual dimorphism at the adult stage can be a consequence of sexual conflict at the larval stage. For example, larval body size and rate of development are likely targets of sexually antagonistic selection (i.e., large size and rapid development are selected for in females but selected against in males). Indeed, genes involved in metabolism, a key determinant of the rate of development in many animals, were enriched in the genes that acquired sex-biased expression on the neo-sex chromosomes at the larval stage. These results indicate that acquiring neo-sex chromosomes may have contributed to a reduction in sexual conflict, particularly at the larval stage, in *Drosophila*. (247/250 words)

## INTRODUCTION

Sex chromosomes are one of the major sex-determination systems and are observed in a variety of organisms, particularly in animals. Considering their evolutionary consequences, including the degeneration of the Y chromosome in many cases, however, we face a difficulty to interpret how the organisms with sex chromosomes have diversified in long-term evolution.

A potential advantage of sex chromosomes is the ability to reduce sexual conflict, in which the optimal phenotype differs between females and males. Under sexual conflict, the optimal phenotype cannot be achieved simultaneously in both sexes (Parker, 2006). Substantial sexual conflict may reduce the fitness of species and potentially lead to extinction. It is estimated that as many as 1,200 genes in *Drosophila melanogaster,* corresponding to ∼8% of the total genes, are under sexual conflict (Innocenti & Morrow, 2010). The accumulation of sexual conflict in a genome is inevitable because sexually antagonistic selection is widespread, even though the genome is largely shared between females and males. Yet, on sex chromosomes, selection can act separately on females and males. This separation is possible because of the female-biased inheritance of the X chromosome (i.e., two-thirds of the X chromosome is inherited through females) and the male-specific inheritance of the Y chromosome. Therefore, theoretically, the acquisition of sex chromosomes can reduce sexual conflict in a species (Rice, 1984, Kasimatis et al., 2017). Indeed, sexually antagonistic alleles are disproportionately found on sex chromosomes (Mank et al., 2014).

If sex chromosomes contribute to the resolution of sexual conflict, species with sex chromosomes are expected to show less sexual conflict than that in species without sex chromosomes. However, this comparison is usually not practical for several reasons. First, measuring and comparing the extent of sexual conflict between species at the phenotypic level is very difficult. Second, species with and without sex chromosomes are often distantly related because the origin of sex chromosomes is ancient, in general [e.g., ∼160 million years ago (Mya) in therian mammals (Veyrunes et al., 2008) and >60 Mya in *Drosophila* (Tamura et al., 2004)]. In this case, it may be difficult to attribute an observed difference in the extent of sexual conflict between species to sex chromosomes, as other differences in the genomes and environmental factors would also contribute.

To overcome the difficulties in empirical analyses of sexual conflict, several approaches are promising. First, instead of directly measuring sexual conflict, sex-biased gene expression can be used as a proxy. This is reasonable because sexual conflict can theoretically be resolved by acquiring sex-biased or sex-specific expression. For example, in *Oryzias woworae*, male-biased expression of *csf1* determines the male-specific red pectoral fin, which contributes to high reproductive success in males and low attractiveness to predatory fish (Ansai et al., 2021). Second, instead of comparing species with and without sex chromosomes, species with and without neo-sex chromosomes can be compared. Neo-sex chromosomes originate when one or both members of a pair of autosomes fuse to either the X or Y chromosome. Therefore, neo-sex chromosomes are, by definition, younger than ordinary sex chromosomes. If sex chromosomes contribute to a reduction of sexual conflict, species with neo-sex chromosomes are expected to acquire sex-biased genes, particularly on the neo-sex chromosomes, after divergence from species without neo-sex chromosomes.

In this study, we evaluated the above hypothesis that acquiring sex-biased gene expression on neo-sex chromosomes reduce sexual conflict using three trios of *Drosophila* species, *D. miranda*–*D. pseudoobscura*–*D. obscura*, *D. albomicans*–*D. nasuta*–*D. kohkoa*, and *D. americana*–*D. texana*–*D. novamexicana*. The first species of each trio possesses neo-sex chromosomes, whereas the second and third species of each trio do not have neo-sex chromosomes. We determined the direction of change in gene expression by using the third species of each trio as an outgroup. Our results provide insight into the evolution of sex-biased gene expression and sexual conflict in *Drosophila* species in relation to the acquisition of neo-sex chromosomes.

## MATERIALS AND METHODS

### Genomes and transcriptomes of three trios of *Drosophila* species

Genome sequences of the nine *Drosophila* species used in this study were retrieved from NCBI (https://www.ncbi.nlm.nih.gov/) (Table S1 for detailed information). RNA-seq data were also downloaded from NCBI (Table S2 for detailed information). Gene annotation data were downloaded from Nozawa et al. (2021) at the journal website (https://genome.cshlp.org/content/31/11/2069/suppl/DC1) as gff files and converted into gtf files.

### Estimation of gene expression

Using the RNA-seq data for the nine species (Nozawa et al., 2016, Nozawa et al., 2021), for each sex, the expression levels of each gene in larvae, pupae, adults, and gonads (testis/ovary) were estimated following the approach described by Nozawa et al. (2021). In short, all qualified read pairs (quality value ≥25 and ≥50 nucleotides for both paired reads) were mapped onto the genome assembly for each species using STAR 2.7 (Dobin et al., 2013) with the gtf file for gene annotation. The mapped data were further processed with RSEM v1.3.1 (Li & Dewey, 2011) to obtain TPM (transcripts per kilobase million mapped reads) (Wagner et al., 2012) for each gene. We also estimated the expression level of each gene in adult heads, thoraxes, abdomens, abdomens without gonads, and accessory glands from *D. miranda*, *D*. *pseudoobscura*, and *D. obscura* as well as in imaginal discs and the remaining larval body carcasses from *D. miranda* and *D. pseudoobscura*.

### Classification of female-biased, male-biased, and unbiased genes

Using the RNA-seq data (Nozawa et al., 2016, Nozawa et al., 2021), we classified each gene in the nine genomes as male-biased, female-biased, or unbiased in each stage or tissue. First, the number of reads mapped to each gene was computed using mmquant v1.3 (Zytnicki, 2017) with the bam file output from STAR. When a read was mapped to multiple genes, it was equally allocated to the genes with our custom Perl script, which is available upon request to MN. Second, for each tissue in each species, genes were classified as male-biased, female-biased, or unbiased using the TCC package in R with a cutoff threshold of *Q* < 0.05 (Sun et al., 2013). Only genes with a 1:1:1 orthologous relationship (see the next section for the identification of orthologs) and located on the same Muller element in the three species of each trio were included in analyses. It should be mentioned that we did not separately treat each gametolog on the neo-X chromosome and the neo-Y chromosome (neo-X and neo-Y, hereafter, respectively) but summed gene expression levels of both gametologs in males with neo-sex chromosomes.

### Identification of orthologs

To identify orthologs in each trio, we followed the procedure described by Nozawa et al. (2021). In short, when a gene showed a relationship of reciprocal best-hit between the outgroup species (e.g., *D. obscura*) and the species with neo-sex chromosomes (e.g., *D. miranda*) and between the outgroup species and the closely-related species without neo-sex chromosomes (e.g., *D. pseudoobscura*) in the BLASTP [ver. 2.9.0+ (Altschul et al., 1997)] search, we regarded the gene as a 1:1:1 ortholog. It should be mentioned that genes on the neo-sex chromosomes were regarded as 1:1:1 orthologs if at least one of the gametologs on the neo-X or the neo-Y was present in the species with neo-sex chromosomes in addition to the genes in the other two species without neo-sex chromosomes.

To identify orthologs among the nine species, we first identified the genes showing a reciprocal best-hit relationship between all combinations of outgroup species in each trio (i.e., *D. obscura*–*D. kohkoa*, *D. obscura*–*D. novamexicana*, and *D. kohkoa*–*D. novamexicana*). If a gene showed a reciprocal best-hit relationship in all three comparisons and was identified as a 1:1:1 ortholog in all three trios, it was regarded as a 1:1:1:1:1:1:1:1:1 ortholog in the nine species. The TPM value and sex-biased status for each gene in each species are listed in Tables S3–5. As mentioned above, raw read counts (and not TPM values) were utilized to identify sex-biased genes.

### Estimation of the proportion of shared sex-biased genes

To compute the proportion (*P*) of shared sex-biased genes at each stage or tissue in each trio, we used the following equation:

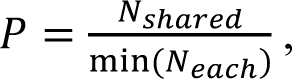

where *N*_shared_ is the number of sex-biased genes shared in all three species and min (*N*_each_) indicates the smallest number of sex-biased genes in the three species. Similarly, the proportion of shared sex-biased genes in the nine species was computed by using the same equation, defining *N*_shared_ as the number of sex-biased genes conserved in all nine species and min (*N*_each_) as the smallest number of sex-biased genes shared in each of the three trios.

## RESULTS

### Sex-biased genes in each species

The number of sex-biased genes increased during development in all nine species (Table 1, Tables S6–S8), consistent with results of previous studies of *Drosophila* (Ingleby & Morrow, 2017, Djordjevic et al., 2022) and other organisms (Ma et al., 2018, Djordjevic et al., 2022). At the larval and pupal stages, there was a general trend that the numbers of sex-biased genes are significantly greater for species with neo-sex chromosomes than species without neo-sex chromosomes in all three trios (Table 1, Tables S6–S8). The greater number of sex-biased genes in species with neo-sex chromosomes than in species without neo-sex chromosomes was generally not confined to the neo-sex chromosomes (in comparison with the homologous autosome in close relatives) but was also observed for other chromosomes. Yet, the trend was more conspicuous on the neo-sex chromosomes in many cases (Table 1). By contrast, in adults and gonads, the differences in the number of sex-biased genes between species with and without neo-sex chromosomes were less clear or there were fewer sex-biased genes in the former than the latter.

**Table 1.**
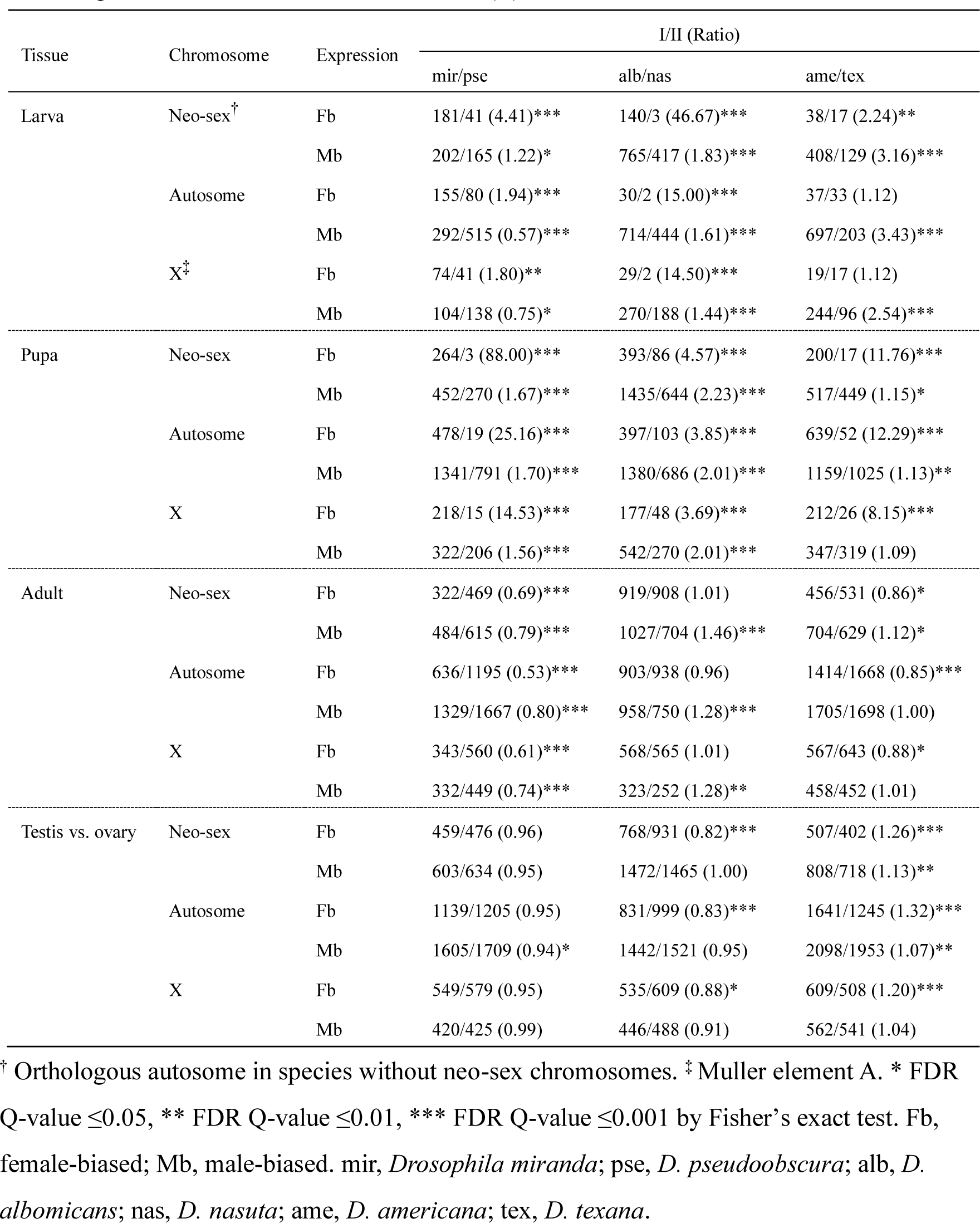
Ratio of the number of sex-biased genes in a species with neo-sex chromosomes (I) to that in a species without neo-sex chromosomes (II).

| Tissue | Chromosome | Expression | I/II (Ratio) |  |  |
| --- | --- | --- | --- | --- | --- |
|  |  |  | mir/pse | alb/nas | ame/tex |
| Larva | Neo-sex<sup>†</sup> | Fb | 181/41 (4.41)*** | 140/3 (46.67)*** | 38/17 (2.24)** |
|  |  | Mb | 202/165 (1.22)* | 765/417 (1.83)*** | 408/129 (3.16)*** |
|  | Autosome | Fb | 155/80 (1.94)*** | 30/2 (15.00)*** | 37/33 (1.12) |
|  |  | Mb | 292/515 (0.57)*** | 714/444 (1.61)*** | 697/203 (3.43)*** |
|  | X<sup>‡</sup> | Fb | 74/41 (1.80)** | 29/2 (14.50)*** | 19/17 (1.12) |
|  |  | Mb | 104/138 (0.75)* | 270/188 (1.44)*** | 244/96 (2.54)*** |
| Pupa | Neo-sex | Fb | 264/3 (88.00)*** | 393/86 (4.57)*** | 200/17 (11.76)*** |
|  |  | Mb | 452/270 (1.67)*** | 1435/644 (2.23)*** | 517/449 (1.15)* |
|  | Autosome | Fb | 478/19 (25.16)*** | 397/103 (3.85)*** | 639/52 (12.29)*** |
|  |  | Mb | 1341/791 (1.70)*** | 1380/686 (2.01)*** | 1159/1025 (1.13)** |
|  | X | Fb | 218/15 (14.53)*** | 177/48 (3.69)*** | 212/26 (8.15)*** |
|  |  | Mb | 322/206 (1.56)*** | 542/270 (2.01)*** | 347/319 (1.09) |
| Adult | Neo-sex | Fb | 322/469 (0.69)*** | 919/908 (1.01) | 456/531 (0.86)* |
|  |  | Mb | 484/615 (0.79)*** | 1027/704 (1.46)*** | 704/629 (1.12)* |
|  | Autosome | Fb | 636/1195 (0.53)*** | 903/938 (0.96) | 1414/1668 (0.85)*** |
|  |  | Mb | 1329/1667 (0.80)*** | 958/750 (1.28)*** | 1705/1698 (1.00) |
|  | X | Fb | 343/560 (0.61)*** | 568/565 (1.01) | 567/643 (0.88)* |
|  |  | Mb | 332/449 (0.74)*** | 323/252 (1.28)** | 458/452 (1.01) |
| Testis vs. ovary | Neo-sex | Fb | 459/476 (0.96) | 768/931 (0.82)*** | 507/402 (1.26)*** |
|  |  | Mb | 603/634 (0.95) | 1472/1465 (1.00) | 808/718 (1.13)** |
|  | Autosome | Fb | 1139/1205 (0.95) | 831/999 (0.83)*** | 1641/1245 (1.32)*** |
|  |  | Mb | 1605/1709 (0.94)* | 1442/1521 (0.95) | 2098/1953 (1.07)** |
|  | X | Fb | 549/579 (0.95) | 535/609 (0.88)* | 609/508 (1.20)*** |
|  |  | Mb | 420/425 (0.99) | 446/488 (0.91) | 562/541 (1.04) |

<sup>†</sup> Orthologous autosome in species without neo-sex chromosomes. <sup>‡</sup> Muller element A. \* FDR Q-value $\leq 0.05$ , \*\* FDR Q-value $\leq 0.01$ , \*\*\* FDR Q-value $\leq 0.001$ by Fisher's exact test. Fb, female-biased; Mb, male-biased. mir, *Drosophila miranda*; pse, *D. pseudoobscura*; alb, *D. albomicans*; nas, *D. nasuta*; ame, *D. americana*; tex, *D. texana*.

We also analyzed RNA-seq data for five adult tissues (i.e., heads, thoraxes, abdomens, abdomens without gonads, and accessory glands/ovaries) in *D. miranda*, *D. pseudoobscura*, and *D. obscura*. In heads and thoraxes, there were more sex-biased genes in *D. miranda* than in *D. pseudoobscura,* particularly male-biased genes, and the trend was more conspicuous on the neo-sex chromosomes than on autosomes and the X chromosome (Table 2, Table S9). The same trend was observed in abdomens after removing gonads; however, the differences were less substantial. We did not observe similar patterns in the comparison of accessory glands and ovaries. Furthermore, abdomens including gonads did not show a trend specific to the neo-sex chromosomes, although the ratio of the number of sex-biased genes in *D. miranda* to that in *D. pseudoobscura* was significantly greater than 1. Collectively, the numbers of sex-biased genes on the neo-sex chromosomes were greater in larvae, pupae, and adult somatic tissues in species with neo-sex chromosomes than in species without neo-sex chromosomes.

**Table 2.** Ratio of sex-biased genes in five adult tissues in *Drosophila miranda* (mir) to *D. pseudoobscura* (pse).

| Tissue | Chromosome | Expression | mir/pse (Ratio) |
| --- | --- | --- | --- |
| Head | Neo-sex<sup>†</sup> | Fb | 52/17 (3.06)*** |
|  |  | Mb | 84/0 (n/a)*** |
|  | Autosome | Fb | 44/24 (1.83)* |
|  |  | Mb | 31/2 (15.50)*** |
|  | X<sup>‡</sup> | Fb | 15/14 (1.07) |
|  |  | Mb | 17/4 (4.25)* |
| Thorax | Neo-sex | Fb | 27/30 (0.90) |
|  |  | Mb | 56/9 (6.22)*** |
|  | Autosome | Fb | 39/75 (0.52)** |
|  |  | Mb | 14/12 (1.17) |
|  | X | Fb | 16/42 (0.38)** |
|  |  | Mb | 8/8 (1.00) |
| Abdomen | Neo-sex | Fb | 338/240 (1.41)*** |
|  |  | Mb | 484/357 (1.36)*** |
|  | Autosome | Fb | 714/581 (1.23)*** |
|  |  | Mb | 1327/1036 (1.28)*** |
|  | X | Fb | 328/244 (1.34)*** |
|  |  | Mb | 360/280 (1.29)*** |
| Abdomen wo gonads | Neo-sex | Fb | 90/28 (3.21)*** |
|  |  | Mb | 155/53 (2.92)*** |
|  | Autosome | Fb | 99/92 (1.08) |
|  |  | Mb | 151/145 (1.04) |
|  | X | Fb | 50/44 (1.14) |
|  |  | Mb | 25/30 (0.83) |
| Accessory gland vs. ovary | Neo-sex | Fb | 369/425 (0.87)* |
|  |  | Mb | 477/512 (0.93) |
|  | Autosome | Fb | 1021/1086 (0.94) |
|  |  | Mb | 1180/1368 (0.86)*** |
|  | X | Fb | 432/431 (1.00) |
|  |  | Mb | 340/417 (0.82)** |

<sup>†</sup> Orthologous autosome in species without neo-sex chromosomes. <sup>‡</sup> Muller element A. \* FDR Q-value $\leq 0.05$ , \*\* FDR Q-value $\leq 0.01$ , \*\*\* FDR Q-value $\leq 0.001$ . Fb, female-biased; Mb, male-biased.

We also computed the proportion of sex-biased genes on the neo-sex chromosome normalized by the proportion on autosomes. The proportion was significantly greater than 1 for both female- and male-biased genes at the larval stage except for male-biased genes in *D. albomicans,* for which the value was close to 1 (Figure 1). However, the proportion decreased with development and was not significantly different from 1 in the gonads except for male-biased genes in *D. americana*. In many cases, such pattern was not observed on orthologous autosomes in the closely related species without neo-sex chromosomes (Figure S1).

**Figure 1.**
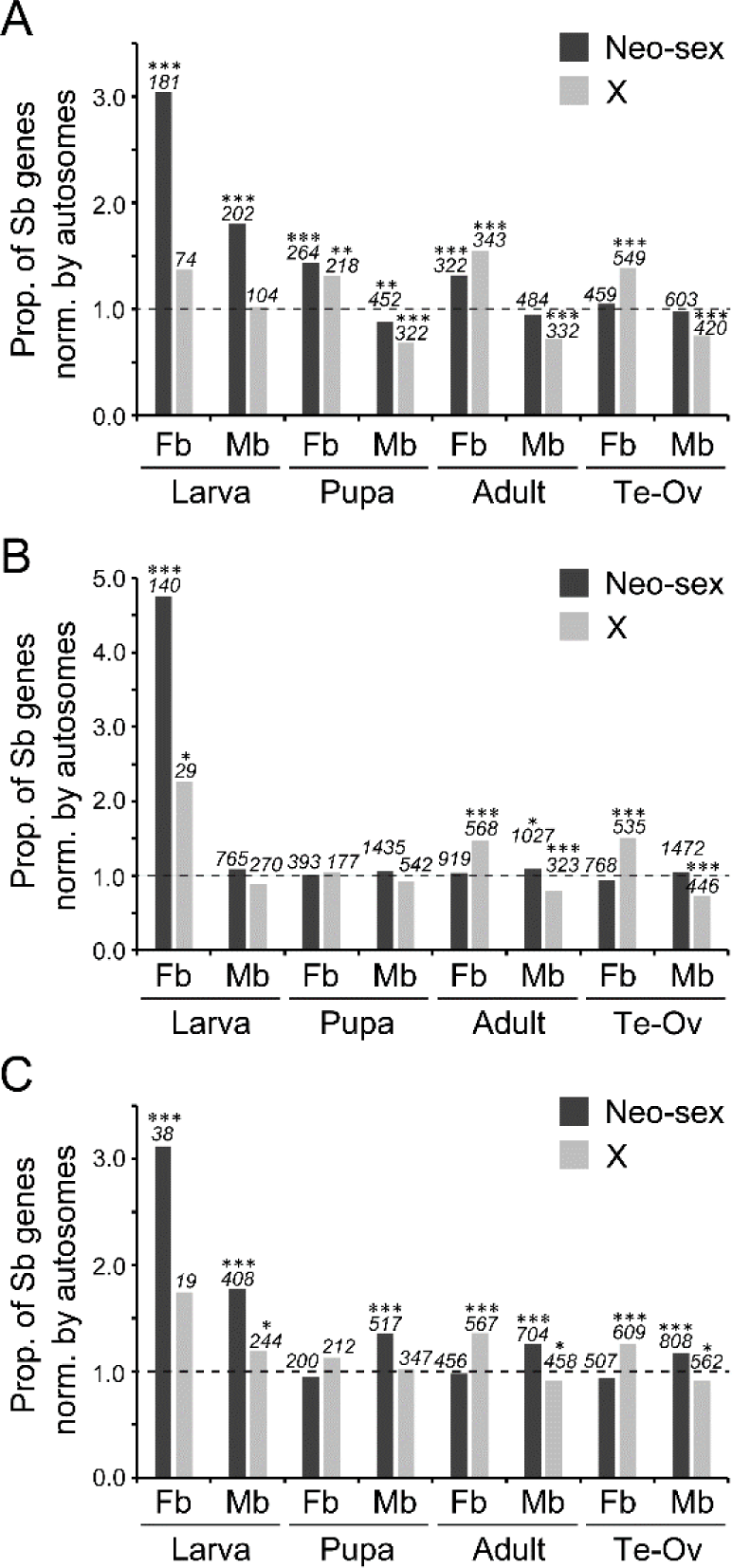
Proportion of sex-biased genes on the neo-sex chromosomes and the X chromosome in (A) *Drosophila miranda*, (B) *D. albomicans*, and (C) *D. americana* normalized by the proportion of sex-biased genes on autosomes. A value of 1.0 shown by a broken line indicates that the proportion of sex-biased genes is equal on the neo-sex (or the X) chromosome and autosomes. The numbers of sex-biased genes on the neo-sex or the X chromosome are indicated in italics above each bar. Differences between the proportion of sex-biased genes on the neo-sex (or the X) chromosome and autosomes were tested by the Fisher’s exact test with correction for multiple testing by the Benjamini-Hochberg method (Benjamini & Hochberg, 1995). *** *Q* <0.001; ** *Q* <0.01; * *Q* <0.05. Sb, sex-biased (either female-or male-biased); Fb, female-biased; Mb, male-biased; Te, testis; Ov, ovary. See Figure S1 for the results for their closely related species.

On the X chromosome, female-biased genes are overrepresented, whereas male-biased genes tend to be underrepresented in many organisms, such as *Drosophila* (e.g., Parisi et al., 2003, Ranz et al., 2003, Meisel et al., 2012) and mice (Reinius et al., 2012). In adults and gonads, we confirmed this trend in all nine species examined in this study (Figure 1, Figure S1). At the pupal stage, the same trend was observed in some species but not in other species. At the larval stage, the trend observed in the adult and gonadal tissues largely disappeared from the X chromosome. These observations indicate that the overrepresentation of female-biased genes and the underrepresentation of male-biased genes on the X chromosome primarily hold at the adult stage, possibly due to the large sexual dimorphism, providing a basis for sexually antagonistic or sex-specific selection at the gene expression level.

### Shared sex-biased genes among closely related species

We next examined the conservation of sex-biased expression among each of the three trios. At the larval and pupal stages, female-biased genes were largely species-specific and the proportion of shared female-biased genes in each trio was very low (Figure 2A–C). (Note that the proportion of shared female-biased genes was 1.0 at the larval stage in the trio of *D*. *miranda*–*D. pseudoobscura*–*D. obscura*; however, there was only one female-biased gene in *D. obscura*.) At the adult stage and in the gonads, however, the number of shared female-biased genes increased drastically, and the proportion was comparable to that of male-biased genes. By contrast, many male-biased genes were shared among each trio throughout development. These patterns imply that sexual dimorphism derived from sex-biased gene expression in adults and gonads is largely conserved among closely related species in *Drosophila*, whereas female-biased genes are largely species-specific in larvae.

**Figure 2.**
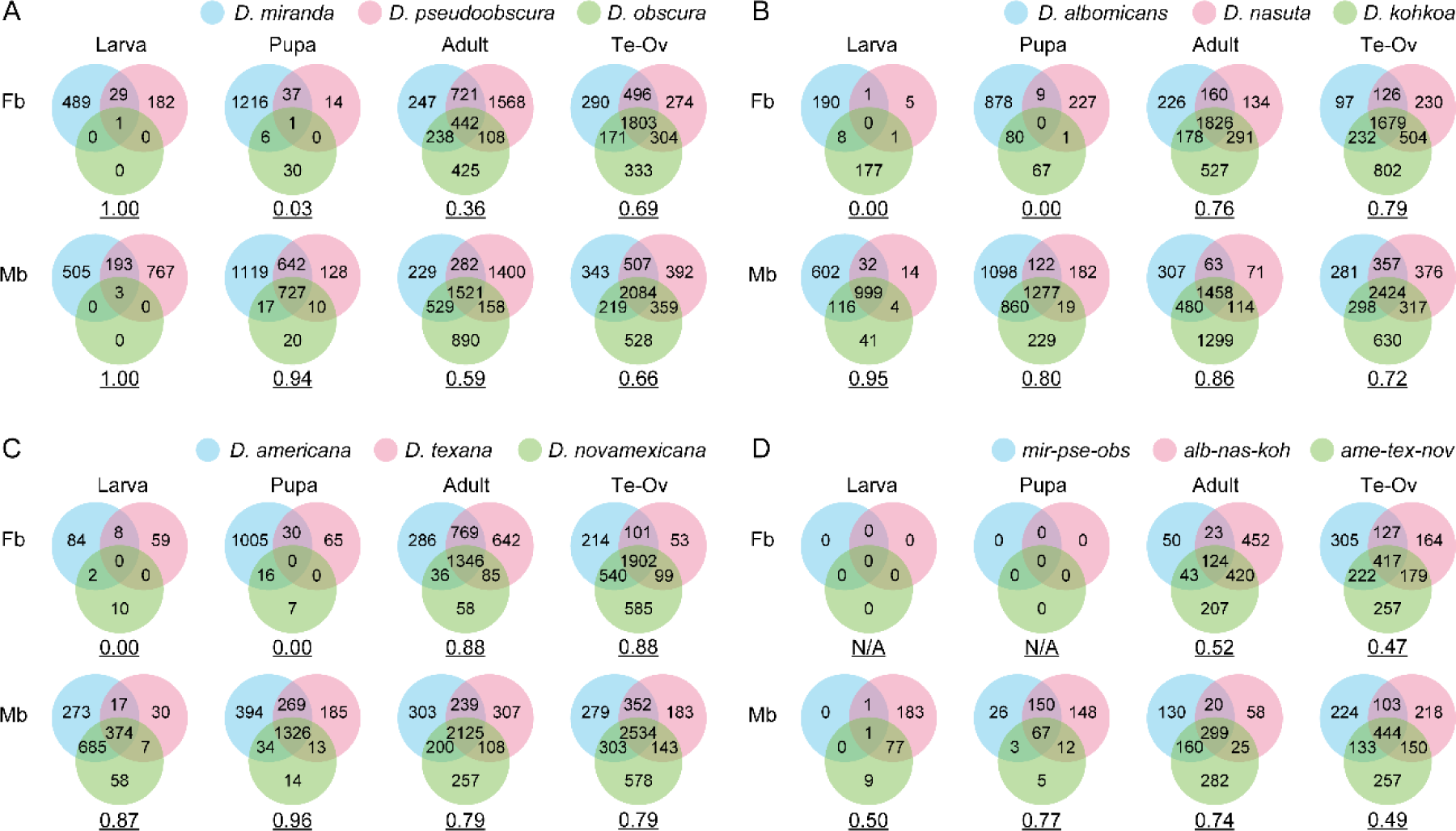
Conservation of sex-biased genes among (A) the trio of *D. miranda*–*D. pseudoobscura*–*D. obscura*, (B) the trio of *D. albomicans*–*D. nasuta*–*D. kohkoa*, (C) the trio of *D. americana*–*D. texana*–*D. novamexicana*, and (D) the three trios. The underlined numbers under each Venn diagram indicate the proportion of shared sex-biased genes among species of interest (see *MATERIALS AND METHODS* for details). In (D), only the genes that were commonly female-biased or male-biased in each trio were analyzed. The number of genes analyzed were (A) 7,804, (B) 8,961, (C) 8,350, and (D) 4,005, respectively. Fb, female-biased; Mb, male-biased; Te, testis; Ov, ovary; *mir*, *D. miranda*; *pse*, *D. pseudoobscura*; *obs*, *D. obscura*; *alb*, *D. albomicans*; *nas*, *D. nasuta*; *koh*, *D. kohkoa*; *ame*, *D. americana*; *tex*, *D. texana*; *nov*, *D. novamexicana*.

We also examined the conservation of sex-biased genes among all nine species. These nine species included two major subgenera, *Sophophora* and *Drosophila*. These subgenera diverged more than 60 Mya (Tamura et al., 2004) but still share quite a few genes with sex-biased expression at the adult stage as well as in gonads (Figure 2D). Consistent with the results for each trio, female-biased genes were not shared at the larval and pupal stages, whereas many male-biased genes were conserved throughout development. The male-biased genes that were shared among nine species tended to be involved in cilium and flagellum movement, male gamete generation, and spermatid differentiation, as determined by an enrichment analysis (Figure S2). By contrast, conserved female-biased genes were associated with many terms related to ribosome and translation, consistent with the necessity of the considerable amount of proteins, such as yolk proteins, required for oogenesis (Hames & Bownes, 1978).

### Lineage-specific sex-biased genes

We detected more sex-biased genes in species with neo-sex chromosomes than in closely related species without neo-sex chromosomes, particularly in larvae. Yet, it is possible that the number of sex-biased genes decreased in species without neo-sex chromosomes rather than increased in species with neo-sex chromosomes after speciation. To test this possibility, using the outgroup species, we estimated the number of genes that acquired sex-biased expression in the lineage with neo-sex chromosomes after splitting from closely related species without neo-sex chromosomes and *vice versa*. For example, in the case of the *D. miranda*–*D. pseudoobscura*–*D. obscura* trio, using *D. obscura* as an outgroup species, unbiased expression in *D. obscura* and *D. pseudoobscura* and male-biased expression in *D. miranda* would indicate that male-biased expression arose in the *D. miranda* lineage after splitting from *D. pseudoobscura* under the parsimony principle.

The number of genes that acquired sex-biased expression after speciation was greater in the species with neo-sex chromosomes than in their close relatives without neo-sex chromosomes in all three trios (Tables S10–S12). However, the pattern was observed only in larvae and pupae but not in adults or gonads. Because this tendency in larvae and pupae was particularly conspicuous for the neo-sex chromosomes (e.g., Muller C in *D. miranda*, Muller C and D in *D. albomicans*, and Muller B in *D. americana*), we calculated the ratio of the proportion of species-specific sex-biased genes on the neo-sex chromosomes to that on autosomes (Figure 3). The ratio was significantly greater than 1 for both female-biased and male-biased genes at the larval stage in all three species with neo-sex chromosomes. The ratio was less conspicuous for other stages and tissues but was still mostly greater than 1. By contrast, the ratio was not significantly greater than 1 in their closely related species except for male-biased genes at the pupal stage of *D. pseudoobscura*. These results indicate that many genes acquired sex-biased expression after linkage to the neo-sex chromosome, particularly in larvae.

**Figure 3.**
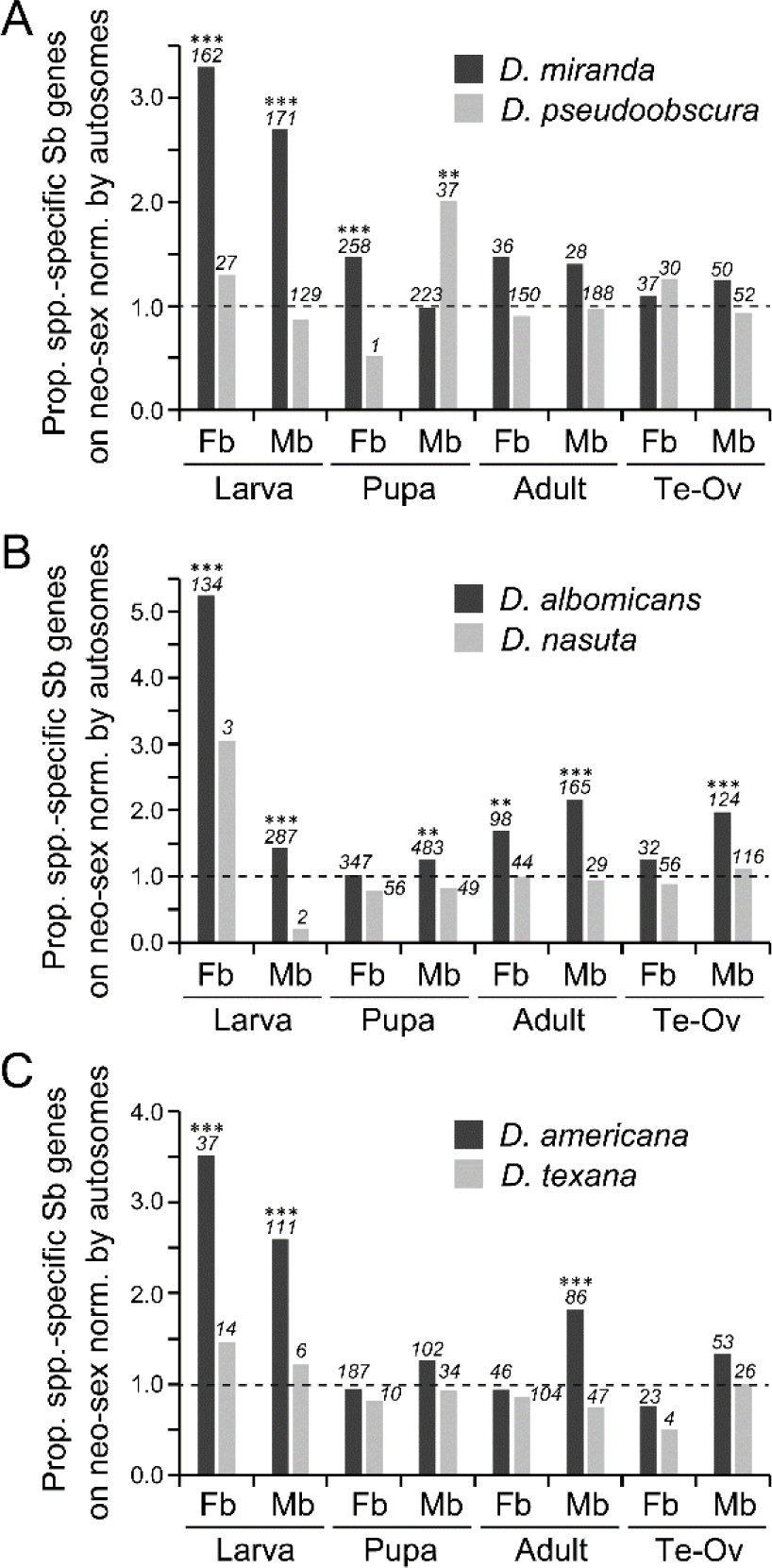
Proportion of genes that acquired sex-biased expression on the neo-sex chromosomes (or the orthologous autosome) in the lineages of (A) *Drosophila miranda* and *D. pseudoobscura*, (B) *D. albomicans* and *D. nasuta*, and (C) *D. americana* and *D. texana* normalized by the proportion of those genes on autosomes. A value of 1.0 by a broken line indicates that the proportion of genes acquiring sex-biased expression is equal in the neo-sex chromosomes (or the orthologous autosome) and autosomes. The numbers of such genes on the neo-sex chromosomes (or the orthologous autosome) are indicated in italics above each bar. Differences between the proportions on the neo-sex chromosomes (or the orthologous autosome) and autosomes were tested by the Fisher’s exact test with correction for multiple testing by the Benjamini-Hochberg method (Benjamini & Hochberg, 1995). *** *Q* <0.001; ** *Q* <0.01; * *Q* <0.05. Sb, sex-biased (either female-or male-biased); Fb, female-biased; Mb, male-biased; Te, testis; Ov, ovary.

To examine the species-specific sex-biased gene expression in adult tissues, we also estimated the number of genes that acquired sex-biased expression after the divergence of *D*. *miranda* and *D. pseudoobscura* by using *D. obscura* as an outgroup species. The proportion of sex-biased genes on the neo-sex chromosomes tended to be significantly higher than that on the autosomes, particularly in heads and thoraxes (Table S13 and Figure 4). By contrast, the orthologous autosome (i.e., chromosome 3) did not show such a pattern in *D. pseudoobscura*. Therefore, in addition to larvae and pupae, adult tissues that do not contain gonads are likely hotspots where genes have acquired sex-biased expression after the birth of neo-sex chromosomes.

**Figure 4.**
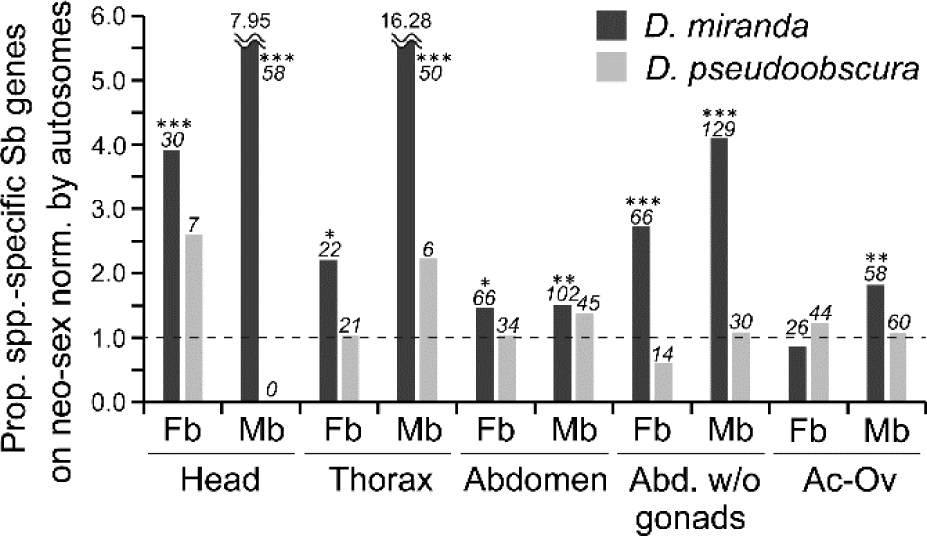
Proportion of genes that acquired sex-biased expression in adult tissues on the neo-sex chromosomes (or the orthologous autosome) in the lineages of *Drosophila miranda* and *D. pseudoobscura*. A value of 1.0 by a broken line indicates that the proportion of genes acquiring sex-biased expression is equal in the neo-sex chromosomes (or the orthologous autosome) and autosomes. The number of such genes on the neo-sex chromosomes (or the orthologous autosome) are indicated in italics above each bar. Differences between the proportions on the neo-sex chromosomes (or the orthologous autosome) and autosomes were tested by the Fisher’s exact test with correction for multiple testing by the Benjamini-Hochberg method (Benjamini & Hochberg, 1995). *** *Q* <0.001; ** *Q* <0.01; * *Q* <0.05. Sb, sex-biased (either female-or male-biased); Fb, female-biased; Mb, male-biased; Te, testis; Ov, ovary.

### Sex-biased gene expression in larval tissues

Our analyses so far revealed that sex-biased gene expression increased on neo-sex chromosomes particularly in larvae. To examine the tissue specificity of sex-biased genes at the larval stage, we also analyzed the RNA-seq data for imaginal discs and other body carcasses separately in *D. miranda* and *D. pseudoobscura* (Nozawa et al., 2016). The number of sex-biased genes was much greater in the body carcass than in imaginal discs (Tables 3 and S14); this result is counter-intuitive because imaginal discs contain genital discs for the testis and ovary, with substantial sexual dimorphism. According to Nozawa et al. (2016), however, the genital discs were not included in the samples of imaginal discs but were included in the samples of other larval body carcasses, explaining these observations.

**Table 3.** Ratio of the number of sex-biased genes in imaginal discs and other larval tissues of *Drosophila miranda* (mir) to that of *D. pseudoobscura* (pse).

| Tissue | Chromosome | Expression | mir/pse (Ratio) |
| --- | --- | --- | --- |
| Imaginal discs | Neo-sex<sup>†</sup> | Fb | 13/3 (4.33)* |
|  |  | Mb | 107/13 (8.23)*** |
|  | Autosome | Fb | 10/11 (0.91) |
|  |  | Mb | 69/31 (2.23)*** |
|  | X<sup>‡</sup> | Fb | 3/12 (0.25) |
|  |  | Mb | 15/15 (1.00) |
| Larval body carcass | Neo-sex | Fb | 105/5 (21.00)*** |
|  |  | Mb | 306/301 (1.02) |
|  | Autosome | Fb | 80/14 (5.71)*** |
|  |  | Mb | 676/849 (0.80)*** |
|  | X | Fb | 33/10 (3.30)*** |
|  |  | Mb | 163/236 (0.69)*** |

<sup>†</sup> Orthologous autosome in species without neo-sex chromosomes. <sup>‡</sup> Muller element A. \* FDR Q-value $\leq 0.05$ , \*\* FDR Q-value $\leq 0.01$ , \*\*\* FDR Q-value $\leq 0.001$ . Fb, female-biased; Mb, male-biased.

The proportion of sex-biased genes also tended to be higher in *D. miranda* with neo-sex chromosomes than in *D. pseudoobscura* without neo-sex chromosomes, particularly on the Muller element C, which became the neo-sex chromosomes in *D. miranda* (Tables 3 and S14). The proportion of species-specific sex-biased genes was also significantly higher on the neo-sex chromosomes than on the autosomes in *D. miranda* but not in *D. pseudoobscura* (Table S15 and Figure 5). In *D. miranda*, the proportion of species-specific sex-biased genes did not differ substantially between the imaginal discs and body carcasses. It should be mentioned that since corresponding data were not available for *D. obscura,* used as an outgroup species, we were unable to exclude the possibility that sex-biased expression has been lost in some genes in the *D. pseudoobscura* lineage after splitting from *D. miranda*. Yet, our analyses consistently indicated that the number of sex-biased genes increased on the neo-sex chromosomes in *D. miranda* rather than decreased on the orthologous autosome in *D. pseudoobscura* at the larval stage.

**Figure 5.**
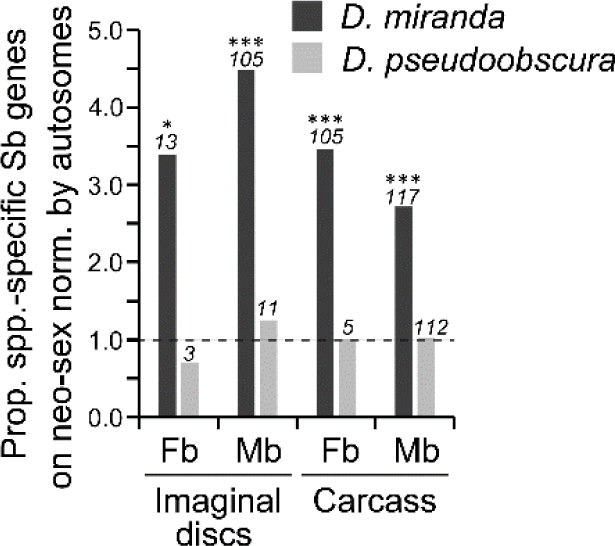
Proportion of genes that acquired sex-biased expression in the larval tissues on the neo-sex chromosomes (or the orthologous autosome) in the lineages of *Drosophila miranda* and *D. pseudoobscura*. A value of 1.0 by a broken line indicates that the proportion of genes acquiring sex-biased expression is equal in the neo-sex chromosomes (or the orthologous autosome) and autosomes. The number of such genes on the neo-sex chromosomes (or the orthologous autosome) is indicated in italics above each bar. Differences between the proportions on the neo-sex chromosomes (or the orthologous autosome) and autosomes were tested by the Fisher’s exact test with correction for multiple testing by the Benjamini-Hochberg method (Benjamini & Hochberg, 1995). *** *Q* <0.001; ** *Q* <0.01; * *Q* <0.05. Sb, sex-biased (either female-or male-biased); Fb, female-biased; Mb, male-biased; Carcass, larval body without imaginal discs.

### Relationship between functional constraints and sex-biased expression

Male-biased genes are known to evolve faster than female-biased and unbiased genes (Meiklejohn et al., 2003). We found the same trend in the three trios (Figure S3). Shared male-biased genes in each trio evolve faster than shared female-biased and shared unbiased genes in terms of nonsynonymous to synonymous substitution rate ratio (*d*_N_/*d*_S_) in the outgroup lineage (i.e., *D. obscura*, *D. kohkoa*, or *D. novamexicana*). This pattern indicates that genes under less functional constraint tend to be male-biased and/or functional constraints became weaker after acquiring male-biased expression. It should be mentioned that the difference in *d*_N_/*d*_S_ between shared male-biased genes and shared unbiased genes was greatest in larvae and decreased with development. In the gonadal tissues, the difference was less clear.

Species-specific male-biased genes in *D. pseudoobscura*, *D. nasuta*, and *D. texana* without neo-sex chromosomes also tended to evolve fast, particularly at the larval stage (Tables S16–18 and Figure 6). By contrast, this pattern was less clear in *D. miranda*, *D. albomicans*, and *D. americana* with neo-sex chromosomes, even in larvae. These results suggest that genes under less functional constraint tend to become male-biased in species without neo-sex chromosomes, particularly in larvae, whereas genes possibly under sexually antagonistic selection with stringent functional constraints may have also become male-biased in species with neo-sex chromosomes as a result of the acquisition of neo-sex chromosomes.

**Figure 6.**
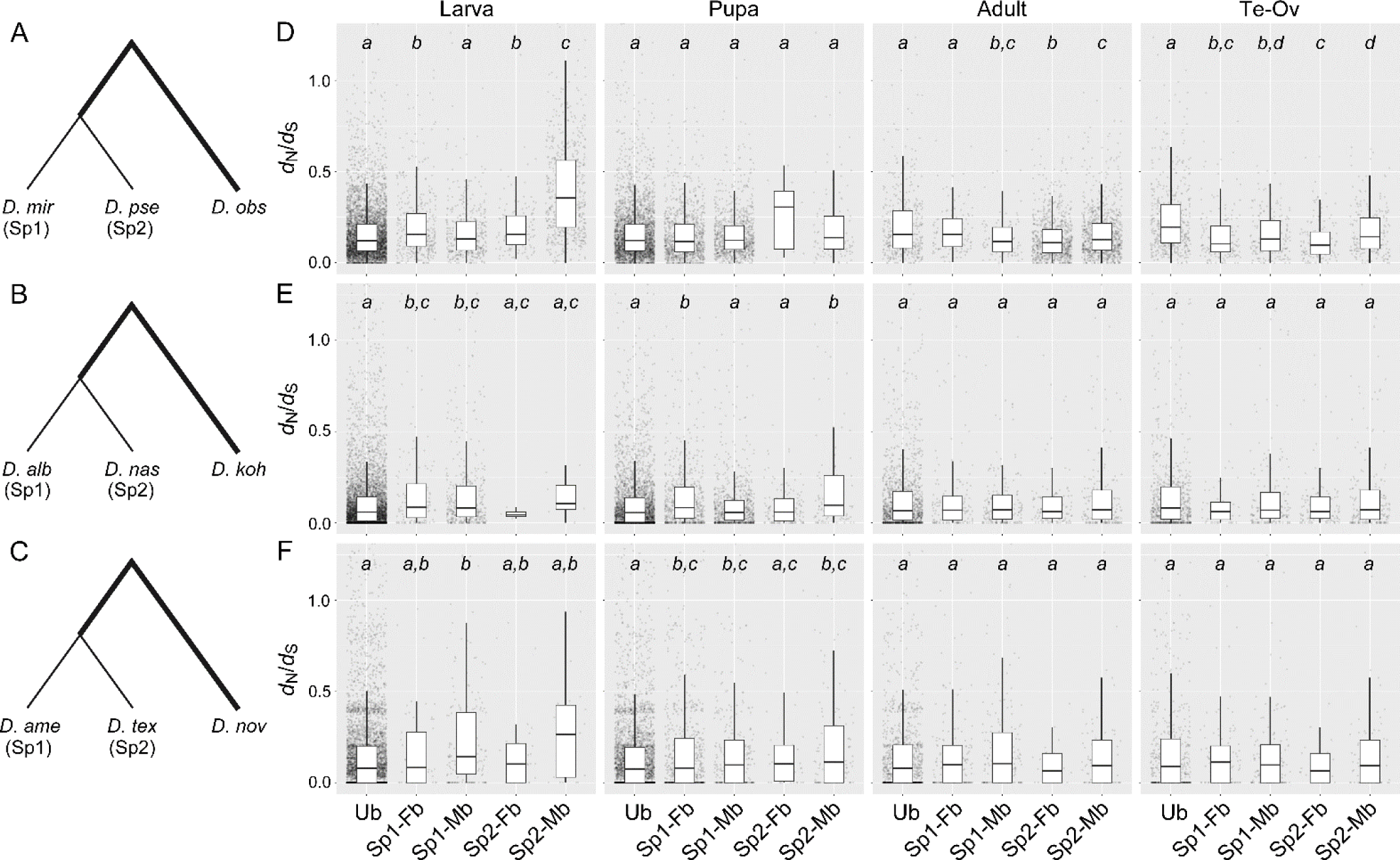
Relationship between functional constraint and sex-biased expression in the lineages of (A and D) *Drosophila miranda* and *D. pseudoobscura*, (B and E) *D. albomicans* and *D. nasuta*, and (C and F) *D. americana* and *D. texana*. The ratio of the numbers of nonsynonymous to synonymous substitutions per site was computed in the lineages of (A) *D. obscura*, (B) *D. kohkoa*, and (C) *D. novamexicana,* as shown in bold, and plotted with five categories for sex-biased expression in larvae, pupae, adults, and gonads (D–F). Ub, unbiased in the trio (e.g., *D. miranda*, *D. pseudoobscura*, and *D. obscura* in D); Sp1-Fb, female-biased only in species 1 (e.g., *D. miranda* in D); Sp1-Mb, male-biased only in species 1; Sp2- Fb, female-biased only in species 2 (e.g., *D. pseudoobscura* in D); Sp2-Mb, male-biased only in species 2. Differences between categories were tested by a Mann–Whitney *U* test with correction for multiple testing by the Benjamini-Hochberg method (Benjamini & Hochberg, 1995). The same letter in italics indicates *Q* ≥ 0.05, whereas different letters indicate *Q* < 0.05 for comparisons between categories.

## DISCUSSION

### Biological significance of acquiring sex-biased genes on neo-sex chromosomes in larvae

We expected the proportion of sex-biased genes in the genome to be higher in species with neo-sex chromosomes than in the species without neo-sex chromosomes because the proportion occupied by the sex chromosomes in a genome is higher in the former than the latter. In particular, we expected the difference to be greater in tissues with marked sexual dimorphism such as gonads (Parisi et al., 2004). However, we did not detect substantial differences in the proportion of sex-biased genes in gonads of the three species that independently acquired neo-sex chromosomes. In addition, sex-biased genes were largely shared among the closely related species in gonads. The conservation of sex-biased gene expression has been reported in four *Drosophila* species (Whittle & Extavour, 2019) as well as in two parasitic wasps (Wang et al., 2015). It is therefore plausible that the number of sex-biased genes already reached a plateau in gonads of the ancestral species before the acquisition of neo-sex chromosomes.

By contrast, we found that many genes acquired sex-biased expression on the neo-sex chromosomes, particularly in larvae, of the three species that independently acquired neo-sex chromosomes. Acquiring sex-biased gene expression can potentially reduce sexual conflict (Parsch & Ellegren, 2013, Ingleby et al., 2015). However, the particularly substantial increase in the number of sex-biased genes in larvae after the gain of neo-sex chromosomes is counter-intuitive because sexual dimorphism is known to increase with development and thus is less conspicuous in larvae, not only at the phenotypic level but also at the transcriptomic (Djordjevic et al., 2022) and metabolomic (Ingleby & Morrow, 2017) levels. Chippindale et al. (2001) also reported that sexual conflict for survival is likely low at the larval stage in *D. melanogaster*.

However, sexual dimorphism at the adult stage may be a consequence of sexual conflict at the larval stage. For example, sexual size dimorphism has frequently been discussed in the context of sexual conflict. There is a positive correlation between the fecundity and body size of females in *D. melanogaster* (Honek, 1993) as well as other insects (Lefranc & Bundgaard, 2000). However, conflicting results regarding the relationship between male body size and fitness have been obtained in *Drosophila*. Several studies of *Drosophila* have reported that larger males are better with respect to maintaining territories (Hoffmann, 1987) and mating success (Partridge & Farquhar, 1983). Yet, crowding is often employed to obtain small males, despite various negative effects, such as low nutrition and high stress. Therefore, the results of these studies should be interpreted with caution. Indeed, several studies have shown that small males are favorable. For example, Pitnick (1991) reported that fecundity is higher in small males than large males in *D. melanogaster*. Using a male-limited experimental evolution approach, Abbott et al. (2010) also reported that smaller males with slower development have a fitness advantage. Because adult flies have an exoskeleton, larval body size is a major determinant of adult body size (Testa et al., 2013). Given the above information, sexual conflict for at least adult body size can likely be attributed to conflict at the larval stage.

To further explore this point, we conducted an enrichment analysis of genes that acquired female-biased or male-biased expression on the neo-sex chromosomes at the larval stage in *D. miranda*, *D. albomicans*, and *D. americana* (Figure S4). We found that many of these genes were enriched for terms related to metabolism, although there were no common terms among the three species with neo-sex chromosomes. Since metabolism is one of the key determinants of developmental rates in many animals (Diaz-Cuadros et al., 2023), some of these genes may have acquired sex-biased expression after linkage to the neo-sex chromosomes and have consequently been released from sexual conflict. We also identified many genes that acquired sex-biased expression on the neo-sex chromosomes in pupae and adult somatic tissues, in which sexual dimorphism is less conspicuous. Therefore, it is possible that there is cryptic sexual conflict at the transcriptional level at the preadult stages and in adult somatic tissues.

### Evolutionary transition from neo-sex chromosomes to ordinary sex chromosomes

In this study, we found that female-biased genes are overrepresented on the ordinary X chromosomes, particularly in adults and gonads, whereas both female- and male-biased genes are enriched on the neo-sex chromosomes in larvae (Figure 1). Female-biased genes were particularly frequent on the neo-sex chromosomes throughout development in *D. miranda*. This is primarily because many neo-Y-linked genes have already become pseudogenes in *D. miranda* and dosage compensation in response to the pseudogenization of neo-Y-linked genes is incomplete on the neo-X (Nozawa et al., 2018, Nozawa et al., 2021); therefore, the expression levels of neo-X-linked genes would become lower in males than in females. In *D. albomicans* and *D. americana*, a strong trend was not detected, possibly because neo-Y degeneration has not progressed substantially owing to the young age of the neo-Y. It should be mentioned that male-biased genes were also overrepresented on the neo-sex chromosomes. If sex-biased selection operates effectively on both the neo-X and neo-Y, many male-biased genes would be primarily transcribed from the neo-Y rather than the neo-X. However, we need to interpret this result with caution because in this study we did not distinguish between the expression of neo-X- and neo-Y-linked gametologs. This is because the genes on the neo-X and neo-Y are mostly identical at the nucleotide sequence level, particularly in *D. albomicans* and *D. americana,* and if only uniquely mapped reads on the neo-X or the neo-Y were used, expression levels of neo-sex-linked genes would be considerably underestimated (Nozawa et al., 2021).

How does the evolutionary transition from young sex chromosomes like neo-sex chromosomes in the three species to ordinary sex chromosomes occur? In *Drosophila*, the ordinary X chromosome originated from an autosome, Muller element A (Vicoso & Bachtrog, 2015), whereas the ordinary Y chromosome is proposed to have been derived from a B chromosome (Hackstein et al., 1996, Carvalho, 2002). Therefore, the evolutionary trajectory of the neo-Y that is homologous to the neo-X would be different from that of the ordinary Y chromosome in *Drosophila* but possibly rather similar to that of the mammalian Y chromosome, although meiotic recombination is absent in males of many *Drosophila* species.

The neo-X contained a higher proportion of female-biased genes in larvae than that on the ordinary X chromosome (Figure 1). As discussed, the large number of female-biased genes on the neo-X in larvae has several explanations, including release from sexual conflict and incomplete dosage compensation. Since a study of the older X chromosome in *D. pseudoobscura* (∼15 Mya) reported more stringent dosage compensation in larvae than in adults (Nozawa et al., 2014), the number of female-biased genes on the neo-X is likely on the way for decreasing in larvae. In this way, the neo-X may exhibit more similar gene expression patterns to those on the ordinary X chromosome in *Drosophila*. Yet, unlike the ordinary X, which emerged through sex-chromosome turnover (Vicoso & Bachtrog, 2013), the neo-X is an additional X chromosome obtained by the fusion of an autosome to the ordinary X and/or Y. Therefore, the evolutionary trajectory of the neo-X may differ from that of the ordinary X chromosome.

### Evolution of heteromorphic sex chromosomes

Our study also provides insights into the process by which X and Y chromosomes diverge. In many organisms, meiotic recombination between X and Y (or Z and W) chromosomes is suppressed except in pseudoautosomal regions, where homologous genes are still present on both chromosomes (Raudsepp & Chowdhary, 2015). However, it is unclear why recombination is suppressed (Wright et al., 2016). According to a widely accepted theory, linkage between a male-determining gene and a male-beneficial gene on the proto-Y chromosome is selected to maintain stable sex determination. Then, recombination is suppressed to maintain stable sex determination and mutations accumulate in this region (Rice, 1996). Recombination suppression in this region can be further strengthened by the accumulation of male-beneficial genes (Rice, 1996). Later, the region of recombination suppression is extended by repeated inversions (Lahn & Page, 1999). In this case, the linkage between the male-determining gene and the male-beneficial gene is the trigger for recombination suppression and the evolution of heteromorphic sex chromosomes.

Theoretically, however, an inversion including the region harboring the male-determining gene can also induce recombination suppression even without the presence of the male-beneficial gene (Wright et al., 2016). We identified many genes that acquired sex-biased expression at the larval stage in the three species with independent neo-sex chromosomes. As discussed above, these genes may have been under sexually antagonistic selection on autosomes but were likely released from such antagonism after being linked to the neo-sex chromosomes. Here, it should be mentioned that many *Drosophila* species, probably including the three species with neo-sex chromosomes, exhibit achiasmy (i.e., no meiotic recombination) in males (John et al., 2016, Charlesworth, 2017), although male recombination likely occurred until recently in *D. albomicans* (Satomura & Tamura, 2016). Therefore, many genes under sexually antagonistic selection (i.e., male-beneficial/female-detrimental genes or *vice versa*) should have accumulated rather quickly after the establishment of recombination suppression. Here, the sexually antagonistic genes are not necessarily a trigger for the establishment of heteromorphic sex chromosomes but just a consequence of recombination suppression. This finding is therefore not inconsistent with the latter possibility that a random inversion including a male-determining gene can generate heteromorphic sex chromosomes in many organisms. A recent simulation study indeed concluded that an inversion including a male-determining gene and later gene-by-gene dosage compensation can establish stable heteromorphic sex chromosomes even without a male-beneficial gene (Lenormand & Roze, 2022).

## CONCLUSION

The acquisition of sex chromosomes can trigger the accumulation of sex-biased genes on sex chromosomes, thereby reducing sexual conflict, particularly at preadult stages. We are currently establishing a method to directly compare the extent of sexual conflict in closely related species with and without neo-sex chromosomes to further verify our conclusion.

## ACKNOWLEDGEMENTS

We thank Shoko Karaki, Takehiro K. Katoh, Jody-Aya Nagasawa, Ayaka Namiki, Yasukazu Okada, Reika Sato, Aya Takahashi, and Koichiro Tamura for their critical comments on earlier versions of the manuscript.

## AUTHOR CONTRIBUTIONS

M.N. and A.M. designed the research; A.M. and M.N. analyzed the data; M.N. and

A.M. wrote the manuscript.

## FUNDING INFORMATION

This work was supported by the Japan Society for the Promotion of Science (JSPS) KAKENHI grant numbers 17H05015, 15K14585, 21H02539, 25711023, 16H06279, and

221S0002 and by Tokyo Metropolitan University.

## DATA ACCESSIBILITY AND BENEFIT-SHARING

Accession numbers of all sequence data used in this study are listed in Tables S1 and S2: https://www.ncbi.nlm.nih.gov/. All sequence data are already deposited by previous studies (Nozawa et al., 2016, Nozawa et al., 2021) which follow the Access to Benefit Sharing.

