## Supplementary figures for "Evolution of sex-biased genes in *Drosophila* species with neo-sex chromosomes: potential contribution to reducing sexual conflict"

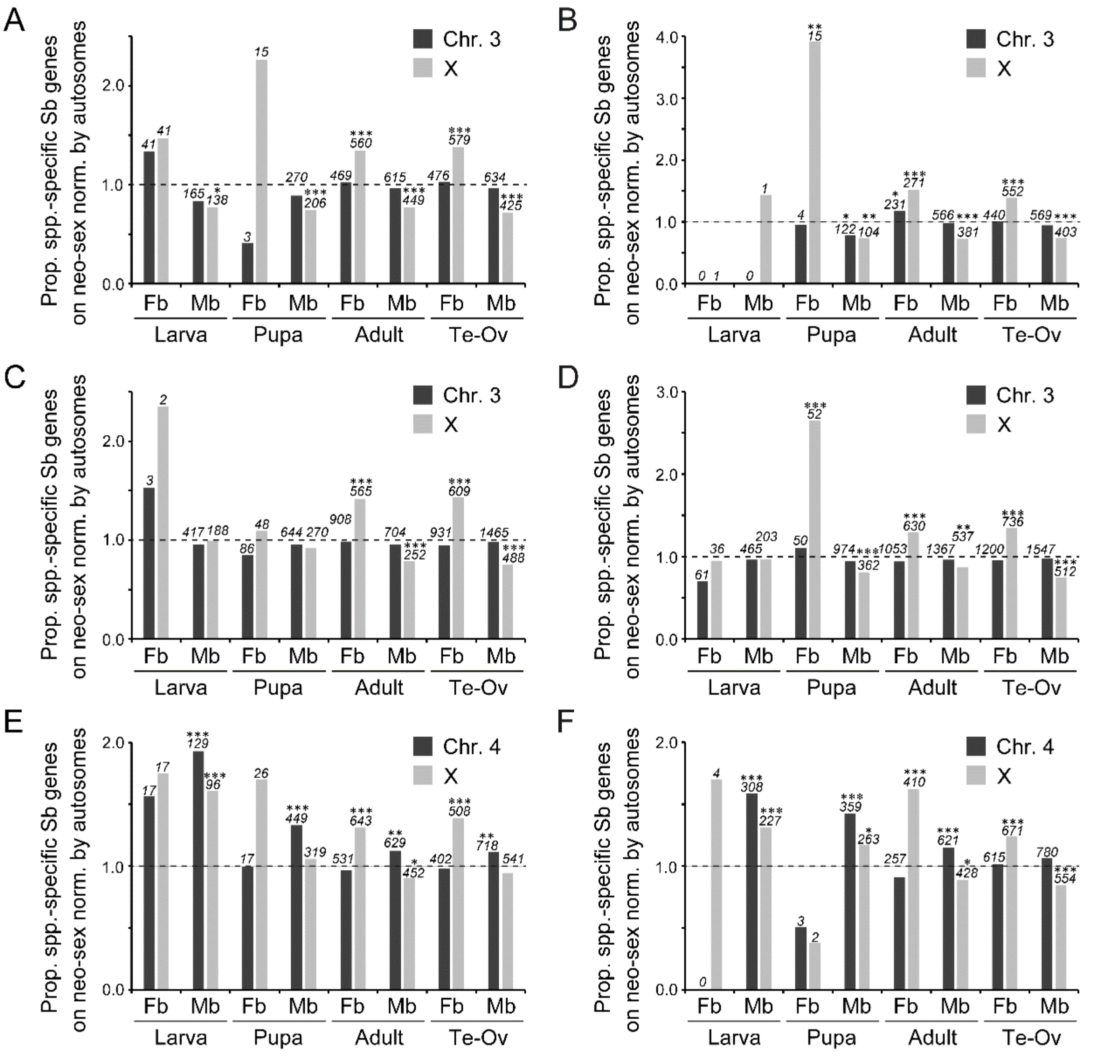


Figure S1. Proportion of sex-biased genes on the autosomes homologous to the neo-sex chromosomes and the X chromosome in (A) *Drosophila pseudoobscura*, (B) *D. obscura*, (C) *D. nasuta*, (D) *D. kohkoa*, (E) *D. texana*, and (F) *D. novamexicana* normalized by the proportion of sex-biased genes on autosomes. A value of 1.0 shown by a broken line indicates that the proportion of sex-biased genes is equal on the target chromosome homologous to the neo-sex chromosomes and autosomes. The numbers of sex-biased genes on the target or the X chromosome are indicated in italics above each bar. Differences between the proportion of sex-biased genes on the target chromosome and autosomes were tested by the Fisher's exact test with correction for multiple testing by the Benjamini-Hochberg method (Benjamini & Hochberg, 1995). *** $Q$ <0.001; ** $Q$ <0.01; * $Q$ <0.05. Sb, sex-biased (either female- or male-biased); Fb, female-biased; Mb, male-biased; Te, testis; Ov, ovary. See Figure S1 for the results for their closely related species.

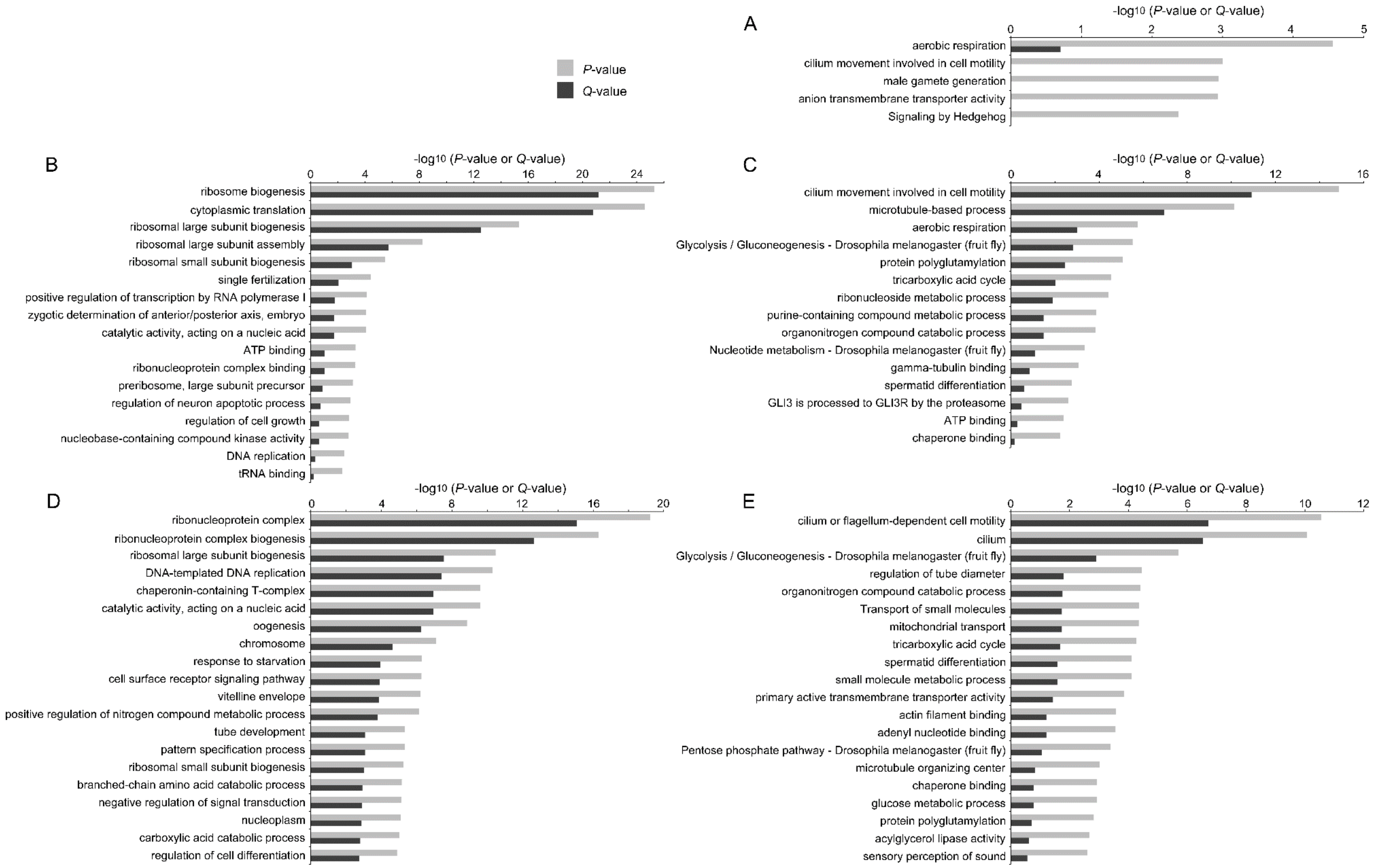
P-value
Q-value
A
-log10 (P-value or Q-value)
0
1
2
3
4
5
aerobic respiration
cilium movement involved in cell motility
male gamete generation
anion transmembrane transporter activity
Signaling by Hedgehog
B
-log10 (P-value or Q-value)
0
4
8
12
16
20
24
ribosome biogenesis
cytoplasmic translation
ribosomal large subunit biogenesis
ribosomal large subunit assembly
ribosomal small subunit biogenesis
single fertilization
positive regulation of transcription by RNA polymerase I
zygotic determination of anterior/posterior axis, embryo
catalytic activity, acting on a nucleic acid
ATP binding
ribonucleoprotein complex binding
preribosome, large subunit precursor
regulation of neuron apoptotic process
regulation of cell growth
nucleobase-containing compound kinase activity
DNA replication
tRNA binding
C
-log10 (P-value or Q-value)
0
4
8
12
16
cilium movement involved in cell motility
microtubule-based process
aerobic respiration
Glycolysis / Gluconeogenesis - Drosophila melanogaster (fruit fly)
protein polyglutamylation
tricarboxylic acid cycle
ribonucleoside metabolic process
purine-containing compound metabolic process
organonitrogen compound catabolic process
Nucleotide metabolism - Drosophila melanogaster (fruit fly)
gamma-tubulin binding
spermatid differentiation
GLI3 is processed to GLI3R by the proteasome
ATP binding
chaperone binding
D
-log10 (P-value or Q-value)
0
4
8
12
16
20
ribonucleoprotein complex
ribonucleoprotein complex biogenesis
ribosomal large subunit biogenesis
DNA-templated DNA replication
chaperonin-containing T-complex
catalytic activity, acting on a nucleic acid
oogenesis
chromosome
response to starvation
cell surface receptor signaling pathway
vitelline envelope
positive regulation of nitrogen compound metabolic process
tube development
pattern specification process
ribosomal small subunit biogenesis
branched-chain amino acid catabolic process
negative regulation of signal transduction
nucleoplasm
carboxylic acid catabolic process
regulation of cell differentiation
E
-log10 (P-value or Q-value)
0
2
4
6
8
10
12
cilium or flagellum-dependent cell motility
cilium
Glycolysis / Gluconeogenesis - Drosophila melanogaster (fruit fly)
regulation of tube diameter
organonitrogen compound catabolic process
Transport of small molecules
mitochondrial transport
tricarboxylic acid cycle
spermatid differentiation
small molecule metabolic process
primary active transmembrane transporter activity
actin filament binding
adenyl nucleotide binding
Pentose phosphate pathway - Drosophila melanogaster (fruit fly)
microtubule organizing center
chaperone binding
glucose metabolic process
protein polyglutamylation
acylglycerol lipase activity
sensory perception of sound

Figure S2. Terms enriched in (A) shared male-biased genes in pupae, (B) shared female-biased genes in adults, (C) shared male-biased genes in adults, (D) shared female-biased genes in gonads, and (E) shared male-biased genes in gonads among the nine *Drosophila* species examined. Only top 20 terms ($P$-value < 0.01) are shown at maximum. Metascape software (Zhou et al. 2019) was used for the analysis.

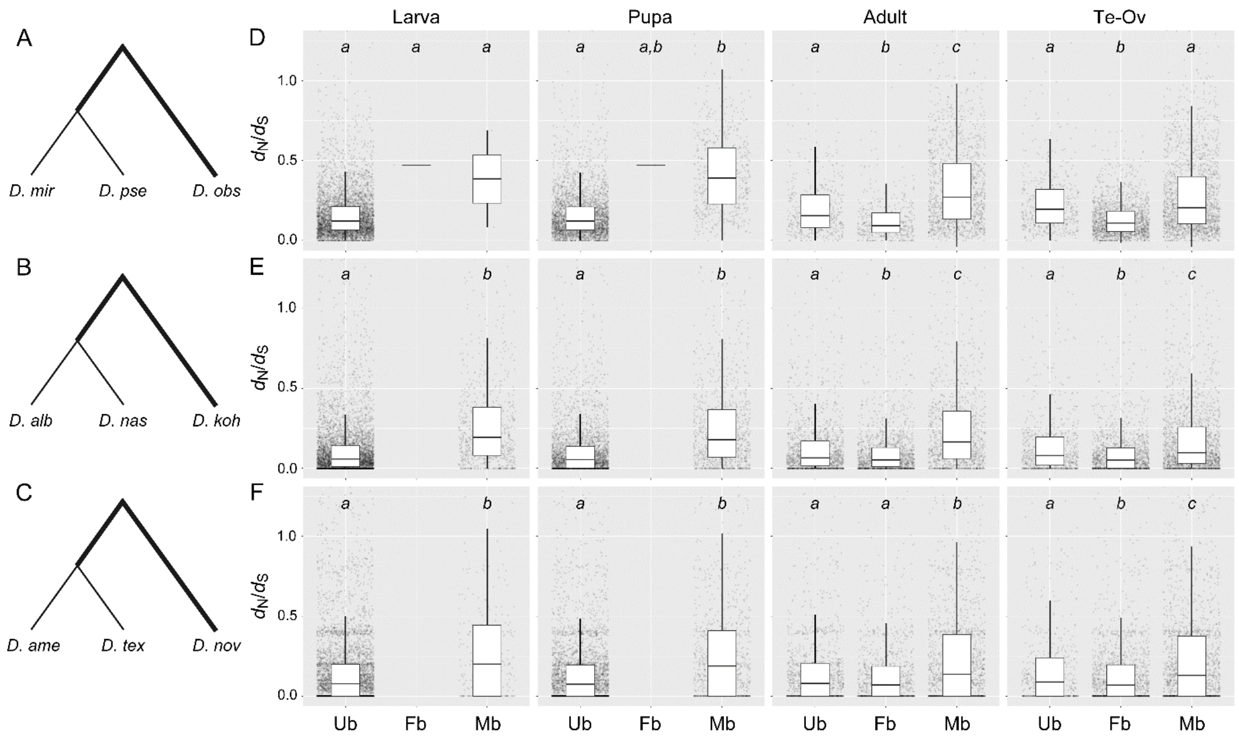


Figure S3. Relationship between functional constraint and sex-biased expression in the lineages of (A and D) *Drosophila miranda* and *D. pseudoobscura*, (B and E) *D. albomicans* and *D. nasuta*, and (C and F) *D. americana* and *D. texana*. The ratio of the numbers nonsynonymous to synonymous substitutions per site was computed in the lineage of (A) *D. obscura*, (B) *D. kohkoa*, and (C) *D. novamexicana* as shown in bold, and plotted with three categories for sex-biased expression in larvae, pupae, adults, and gonads (D-F). Ub, shared unbiased genes in the trio (e.g., *D. miranda*, *D. pseudoobscura*, and *D. obscura* in D); Fb, shared female-biased genes in the trio; Mb, shared male-biased genes in the trio. Differences between categories were tested by a Mann–Whitney *U* test with correction for multiple testing by the Benjamini-Hochberg method (Benjamini & Hochberg, 1995). The same letter in italics indicates $Q \geq 0.05$, whereas different letters indicate $Q < 0.05$ for comparisons between categories.

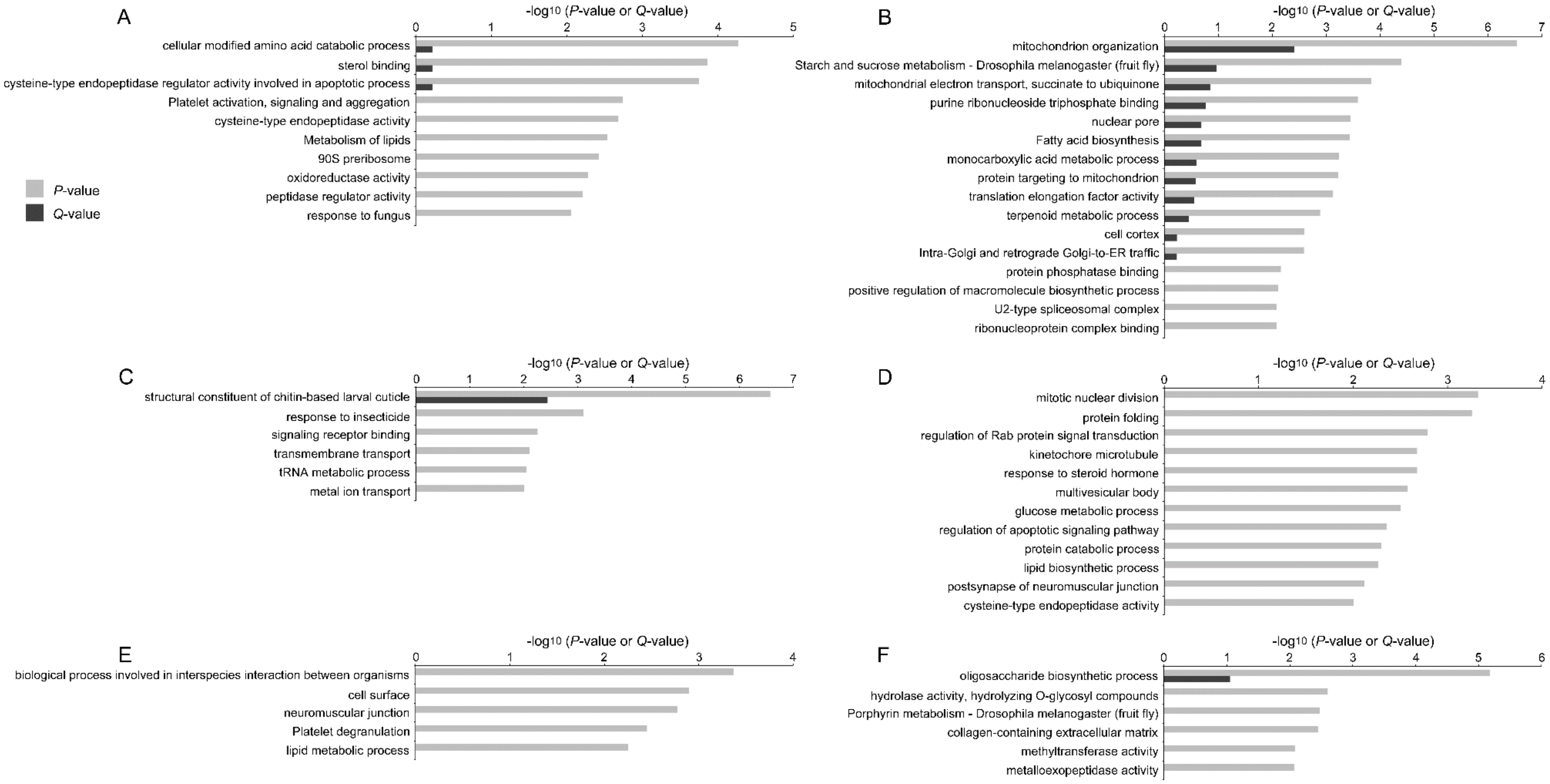


Figure S4. Terms enriched in species-specific sex-biased genes on the neo-sex chromosomes in (A and B) the *Drosophila miranda*, (C and D) the *D. albomicans*, and (E and F) the *D. americana* larvae. The genes are unbiased genes in the other two closely-related species of each trio. (A, C, and E) Species-specific female-biased genes. (B, D, and F) Species-specific male-biased genes. Only top 20 terms (*P*-value < 0.01) are shown at maximum. Metascape software (Zhou et al. 2019) was used for the analysis.
